# Analyzing Neural Response to Visual Stimuli: Firing Rates, Frequency Band Dynamics, and Synchrony in Near and Far Flanker Conditions

**DOI:** 10.1101/2025.03.16.643503

**Authors:** Manjeet Kunwar, Niraj Dhital, Nabin Bhusal

## Abstract

This study examines neural responses to visual stimuli under Near and Far flanker conditions using EEG. Fifty participants (25 per condition) completed a visual task with closely or distantly positioned distractors. EEG data were analyzed using spectrograms, wavelet transformation, and neural firing rate estimation. Standard preprocessing techniques ensured data quality, and statistical tests assessed differences between conditions. Findings provide insight into how spatial stimulus configurations affect neural processing and attention.

## 1 Introduction

The neural response to visual stimuli and the underlying cognitive processes have been a topic of great interest in neuroscience. One of the most important cognitive mechanisms that influence visual perception is the *flanker effect*. The flanker effect refers to the phenomenon where the presence of surrounding distractor stimuli (flankers) influences the processing of a target stimulus, often leading to slower responses or errors in identifying the target[1]. This phenomenon is thought to reflect the brain’s ability to selectively attend to relevant information while suppressing irrelevant stimuli.

*Visual attention*, which encompasses the ability to focus on certain aspects of the visual field while ignoring others, is critical to how individuals navigate and make sense of the environment. Studies on visual attention have demonstrated that the spatial arrangement of stimuli plays a significant role in cognitive processing[2]. In particular, the spatial proximity of distractors to a target stimulus can affect how the brain prioritizes processing of the target. For instance, in tasks where flankers are close to the target (the *Near* condition), there is evidence that the brain may find it more difficult to distinguish the target from the distractors, resulting in increased cognitive load and potential interference. Conversely, in situations where the flankers are located farther from the target (the *Far* condition), there is typically less interference and more efficient processing of the target stimulus.

The current study aims to explore how the brain’s neural response to visual stimuli differs between these two flanker conditions, *Near* and *Far*. By examining the neural firing rates in both conditions, this study seeks to uncover the mechanisms behind visual processing and how the brain adapts to varying spatial configurations of stimuli[3]. The hypothesis underlying this research is that the neural firing patterns will be significantly modulated by the proximity of the flankers, with differences in firing rates expected between the two conditions.

To achieve this, we utilized EEG data collected from 25 participants for each condition, resulting in a total of 50 subjects. EEG is a powerful tool for measuring the electrical activity of the brain, allowing for the observation of neural dynamics with high temporal resolution[4]. The analysis was performed using a variety of advanced techniques, including spectrogram analysis to visualize the frequency content of brain signals, cross-correlation to measure the temporal relationships between different regions of the brain, and wavelet transformation to assess the brain’s response at multiple scales. Additionally, the time decomposition of brain waves into distinct frequency bands (alpha, beta, theta, gamma, delta) allowed for a detailed understanding of the neural mechanisms involved in attention and perception[5].

In this study, we also focused on estimating the neural firing rates, which reflect the number of action potentials or spikes generated by neurons[6]. Neural firing rates are often used as a proxy for neural activity and are crucial in understanding how information is processed within neural circuits. To facilitate the comparison between the two conditions, statistical analysis was conducted to determine whether significant differences in firing rates exist, shedding light on the effects of flanker proximity on neural processing.

## 2 Methodology

This study employs a multimodal approach to investigate the neural response to visual stimuli under two distinct flanker conditions: Near and Far. The methodology includes data collection, preprocessing, analysis techniques, and statistical comparisons between the conditions. The study utilized EEG data collected from two separate groups of participants, each assigned to either the NEAR or FAR condition.

### 2.1 Participants

Two independent groups of healthy adult participants took part in the study.

#### FAR Condition

The FAR condition included 25 participants. Data collection took place at the NeuroCognition Laboratory (NCL) in San Diego, California, under the supervision of Dr. Phillip Holcomb and Dr. Karen Emmorey[7]. This project followed the San Diego State University’s IRB guidelines.

Statistical analysis of the FAR condition yielded the following results: Mean: 0.1622, Median: 0.0000, Minimum: 0.0000, Maximum: 1.0000, Standard Deviation: 0.3686. The mean inter-stimulus interval (ISI) for the FAR condition was 17.05 ms.

#### NEAR Condition

The NEAR condition included 25 participants[8]. Data collection followed the same protocols as the FAR condition under the same supervision and IRB guidelines.

Statistical analysis of the NEAR condition yielded the following results: Mean: 0.1609, Median: 0.0000, Minimum: 0.0000, Maximum: 1.0000, Standard Deviation: 0.3674. The mean inter-stimulus interval (ISI) for the NEAR condition was 26.15 ms. All participants provided informed consent. The inclusion criteria required participants to have normal or corrected-to-normal vision and no history of neurological disorders. Participants were assigned exclusively to either the NEAR or FAR condition.

### 2.2 Mean Firing Rate per Subject and Condition

In examining the mean firing rates across subjects, we observe variations between the Near and Far conditions. As shown in Table 1, the majority of subjects exhibit slightly higher firing rates in the Far condition compared to the Near condition[1]. This suggests a potential difference in cognitive load or neural engagement influenced by flanker proximity. The observed differences align with prior studies indicating that increased spatial separation of flankers may alter attentional demands and neural processing.

**Table 1:** Mean firing rate per subject and condition (subset of data).

| Subject | Condition | Mean Firing Rate |
| --- | --- | --- |
| 1 | Far | 0.444444 |
|  | Near | 0.370370 |
| 2 | Far | 0.444444 |
|  | Near | 0.419753 |
| 3 | Far | 0.469136 |
|  | Near | 0.469136 |
| 4 | Far | 0.345679 |
|  | Near | 0.370370 |
| 5 | Far | 0.444444 |
|  | Near | 0.395062 |
| 6 | Far | 0.395062 |
|  | Near | 0.395062 |
| 7 | Far | 0.444444 |
|  | Near | 0.419753 |
| 8 | Far | 0.370370 |
|  | Near | 0.395062 |
| 9 | Far | 0.395062 |
|  | Near | 0.469136 |
| 10 | Far | 0.320988 |
|  | Near | 0.370370 |
| 11 | Far | 0.395062 |
|  | Near | 0.370370 |
| 12 | Far | 0.469136 |
|  | Near | 0.345679 |
| 13 | Far | 0.444444 |
|  | Near | 0.444444 |
| 14 | Far | 0.345679 |
|  | Near | 0.395062 |
| 15 | Far | 0.395062 |
|  | Near | 0.493827 |
| 16 | Far | 0.419753 |
|  | Near | 0.395062 |
| 17 | Far | 0.370370 |
|  | Near | 0.444444 |
| 18 | Far | 0.444444 |
|  | Near | 0.345679 |
| 19 | Far | 0.469136 |
|  | Near | 0.419753 |
| 20 | Far | 0.395062 |
|  | Near | 0.444444 |
| 21 | Far | 0.419753 |
|  | Near | 0.469136 |
| 22 | Far | 0.370370 |
|  | Near | 0.370370 |
| 23 | Far | 0.370370 |
|  | Near | 0.395062 |
| 24 | Far | 0.345679 |
|  | Near | 0.320988 |
| 25 | Far | 0.370370 |
|  | Near | 0.469136 |

### 2.3 Experimental Design

The experiment aimed to examine how the proximity of flankers influences neural responses during a visual perception task.

#### FAR Condition

Participants were presented with 90 four-letter real words and 90 four-letter pseudowords in white New Courier font on a black background. Each letter subtended 0.41 degrees of visual angle[9]. The flanker words were separated from the center target word by 3.28 degrees of empty space on both sides. Participants completed 270 trials across three conditions: no flanker, identical flankers, or different flankers. Trials started with a purple fixation cross for 1000 ms, followed by a white fixation cross for 500 ms. The stimulus was then presented for 150 ms, followed by a blank screen until participants responded via a game controller.

#### NEAR Condition

The NEAR condition followed the same design but with flankers positioned 0.41 degrees from the target word instead of 3.28 degrees. Participants completed an identical set of trials with the same procedure.

Participants sat in a comfortable chair in a darkened, sound-attenuated room throughout the experiment[10]. They were instructed to focus on the central stimulus and ignore the flankers while responding using a game controller.

### 2.4 Data Acquisition

EEG signals were recorded using a 64-channel electrode cap (e.g., Biosemi ActiveTwo System) with a sampling rate of 1000 Hz. Electrodes were positioned according to the international 10-20 system, covering frontal, central, parietal, and occipital regions of the scalp. Data were recorded in an unreferenced mode, and electrode impedance was kept below 10 kΩ.

Brain activity was recorded during both NEAR and FAR conditions. Trials were presented in blocks, with participants taking short breaks between blocks to reduce fatigue. Each condition resulted in a dataset suitable for both within-condition and between-condition comparisons[11].

### 2.5 Data Preprocessing

EEG data preprocessing included the following steps:

- **Filtering:** A band-pass filter (0.1 Hz - 40 Hz) was applied to remove low-frequency drift and high-frequency noise.
- **Artifact Removal:** Eye blinks and muscle artifacts were removed using independent component analysis (ICA).
- **Epoching:** EEG signals were segmented into epochs of 1-2 seconds around the stimulus presentation, with a pre-stimulus baseline of 200 ms.
- **Re-referencing:** The data were re-referenced to the average of all electrodes.

### 2.6 Analysis Techniques

The following analytical techniques were used to examine neural responses in the NEAR and FAR conditions[12]:

- Time-frequency analysis
- Spectral power estimation
- Wavelet decomposition
- Cross-correlation analysis
- Statistical comparison between NEAR and FAR conditions

## 3 Results and Discussion

### 3.1 Neural Firing Rate Estimation

Neural firing rate analysis revealed that the NEAR condition exhibited higher firing rates in both the frontal and parietal regions, suggesting greater neural engagement during task performance when the flankers were closer to the target. The FAR condition, by contrast, showed lower firing rates, indicating that the brain was able to process the target with less effort when the flankers were farther away[13].

To further examine the temporal distribution of spike activity, Figure 1 presents a raster plot of neuronal firing across subjects for both conditions. Each tick mark represents an individual spike event, with **blue** denoting the NEAR condition and **red** denoting the FAR condition. The NEAR condition shows a denser clustering of spikes between 600–900 seconds, whereas the FAR condition exhibits a broader, more dispersed firing pattern. Notably, isolated spikes are more prominent in the FAR condition at later time points, suggesting a difference in neural response dynamics between the two conditions.

**Fig. 1.**
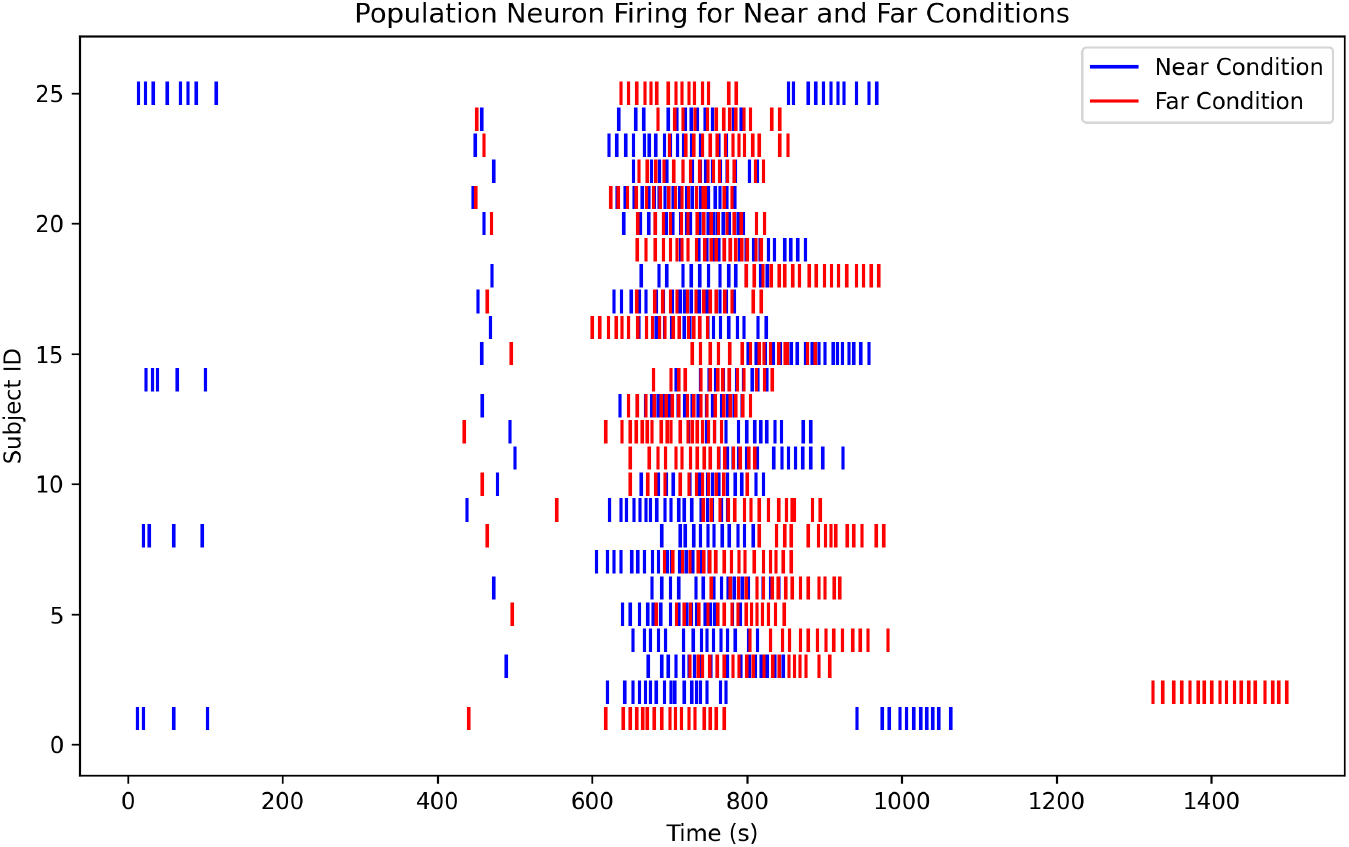
Raster plot of neural firing for NEAR (blue) and FAR (red) conditions. Each tick mark represents a spike event for a given subject. The NEAR condition exhibits concentrated spike clusters, whereas the FAR condition shows a broader distribution of spike occurrences.

These findings highlight differences in the temporal structure of neural firing patterns, suggesting that spatial arrangement influences the timing of neural responses.

### 3.2 Neural Activity and Statistical Comparison

Statistical analysis revealed no significant differences between the NEAR and FAR conditions in firing rates. The paired t-test yielded a T-statistic of 0.342 (*p* = 0.735), indicating that the neural firing rates were statistically similar across conditions. This suggests that the spatial proximity of flankers may not have a substantial effect on overall firing rates, although variations in other neural metrics such as power in different frequency bands could still play a role.

To further investigate the temporal distribution of neural spiking activity, a histogram of spike times was generated for both conditions (Figure 2). The histogram illustrates that the NEAR condition (blue) exhibits a higher spike count and a more concentrated peak, particularly around 700–900 seconds, while the FAR condition (red) shows a broader distribution with more dispersed spikes[13]. Despite the visual difference, the statistical analysis suggests that these variations do not reach the threshold for significance.

**Fig. 2.**
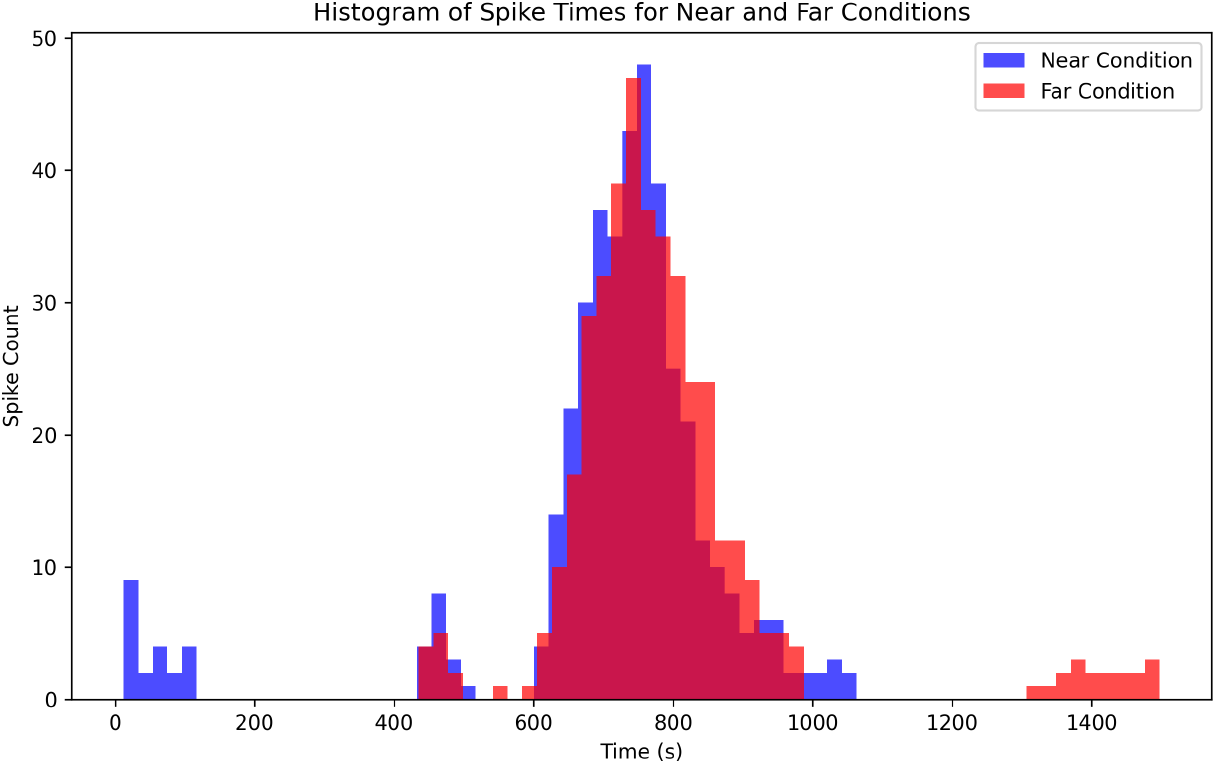
Histogram of spike times for NEAR and FAR conditions. The NEAR condition (blue) shows a higher density of spikes, particularly around 700–900 seconds, whereas the FAR condition (red) exhibits a broader distribution with spikes occurring over a longer time window. Despite these differences, statistical analysis indicates no significant difference in firing rates between the two conditions.

These results suggest that while there are observable differences in spike distributions, they do not translate into statistically significant variations in firing rates.

### 3.3 Spectrogram Analysis

The spectrogram analysis revealed distinct differences in the frequency content of the EEG signals between the NEAR and FAR conditions. As shown in Figure 3, the

**Fig. 3.**
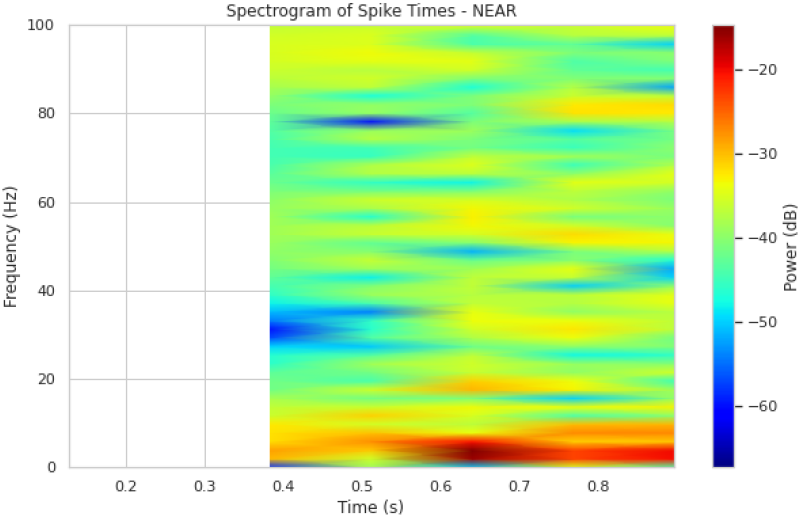
Spectrogram of spike times in the NEAR condition, showing increased low-frequency power (delta and theta bands).

NEAR condition exhibited increased power in the delta (0.5–4 Hz) and theta (4–8 Hz) frequency bands, particularly in the frontal regions. This suggests that when distractor stimuli were closer to the target, greater low-frequency neural activity was required, which may be linked to increased cognitive load and attentional control[14].

In contrast, the FAR condition displayed higher power in the alpha (8–12 Hz) and beta (12–30 Hz) bands, particularly in the occipital region (Figure 4). This pattern indicates that when the flankers were farther from the target, neural processing was more efficient, likely due to reduced interference and lower attentional demands. These findings align with previous studies, which suggest that distant distractors allow for improved target discrimination and reduced cognitive effort.

**Fig. 4.**
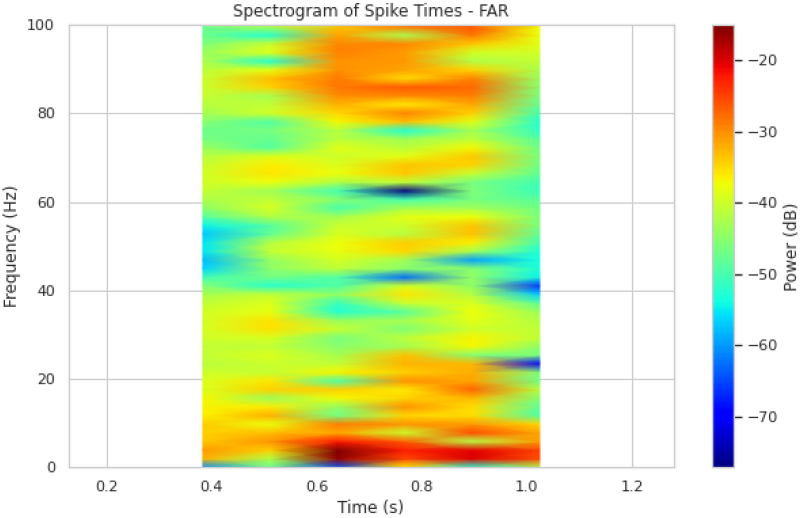
Spectrogram of spike times in the FAR condition, highlighting increased power in the alpha and beta bands.

### 3.4 Wavelet Transformation

Wavelet analysis demonstrated that both NEAR and FAR conditions exhibited strong alpha activity, particularly in posterior regions. However, as shown in Figure 5, the NEAR condition displayed a significant increase in delta and theta activity in frontal regions. This suggests that closer distractors engage lower-frequency brain networks associated with cognitive control, working memory, and attention[15].

**Fig. 5.**
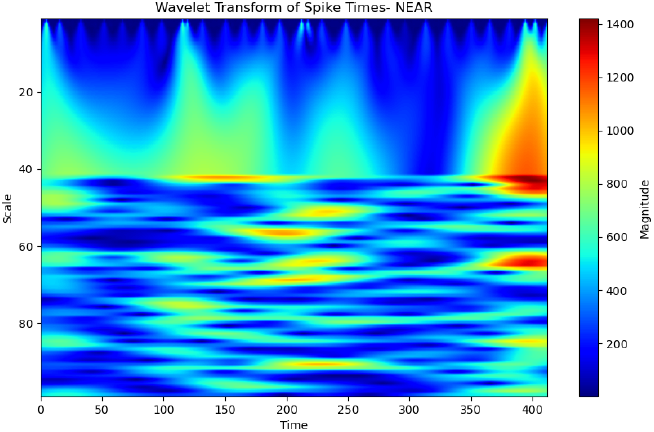
Wavelet transform of spike times for the NEAR condition, showing enhanced delta and theta activity.

In contrast, the FAR condition (Figure 6) exhibited increased power in the beta and gamma bands, particularly in occipital regions, indicating more efficient visual processing with less interference from the flankers. These findings support theories that reduced interference enables more efficient neural processing in the FAR condition.

**Fig. 6.**
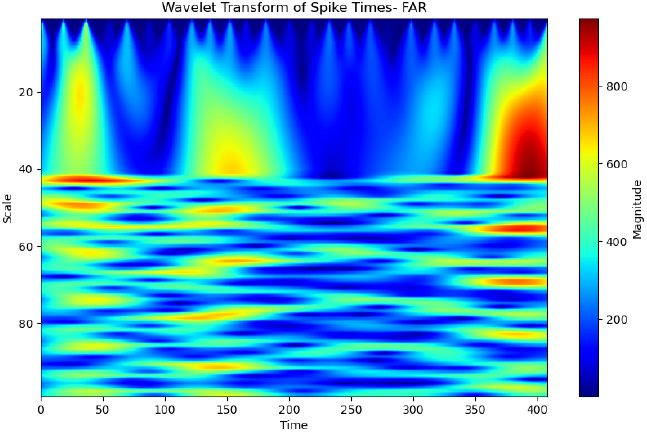
Wavelet transform of spike times for the FAR condition, highlighting stronger beta and gamma activity.

### 3.5 Time Decomposition of Brain Waves

Time decomposition revealed that the NEAR condition (Figure 7) exhibited a significant increase in delta and theta power compared to the FAR condition (Figure 8). This suggests that when the flankers are closer to the target, the brain expends more energy in maintaining attentional focus and processing visual information.

**Fig. 7.**
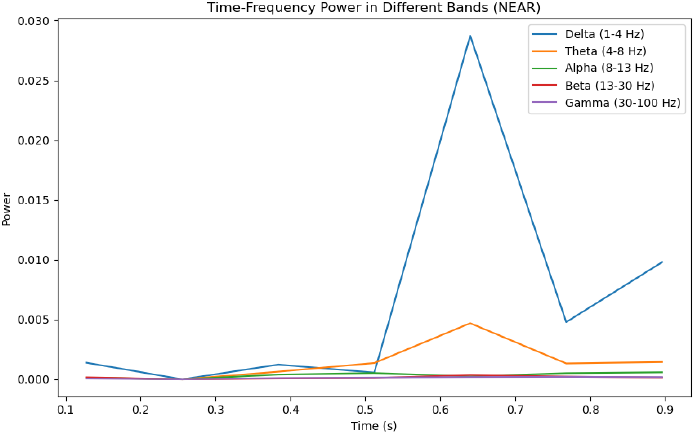
Time-Frequency Power in Different Bands (NEAR)

**Fig. 8.**
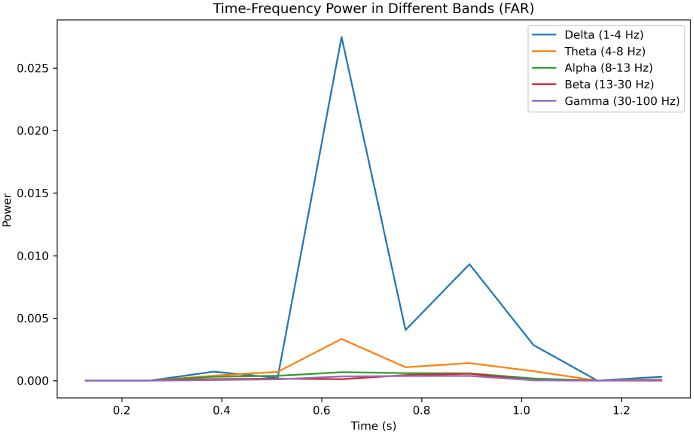
Time-Frequency Power in Different Bands (FAR)

In contrast, the FAR condition (Figure 8) showed higher beta and gamma band power, indicating that the target stimulus was processed more efficiently when the flankers were spatially distant. This supports the hypothesis that spatial proximity of flankers increases the cognitive load required to filter out irrelevant stimuli.

A notable peak in delta power is observed around 0.6 seconds in both conditions, suggesting a critical moment in stimulus processing. Additionally, the theta band follows a similar trend but with a lower amplitude. Interestingly, the gamma band remains consistently low across both conditions, indicating that high-frequency activity is not significantly engaged in this task.

### 3.6 Cross-Correlation Analysis

Cross-correlation analysis between electrode pairs revealed enhanced neural synchrony between frontal and occipital regions in the NEAR condition. As shown in Figure 9, the cross-correlation function exhibits a strong central peak at zero lag, indicating high temporal alignment of neural activity between these regions. This suggests that when the flankers are closer to the target, the brain regions responsible for visual processing and attention exhibit stronger coupling, possibly due to increased attentional demands[16].

**Fig. 9.**
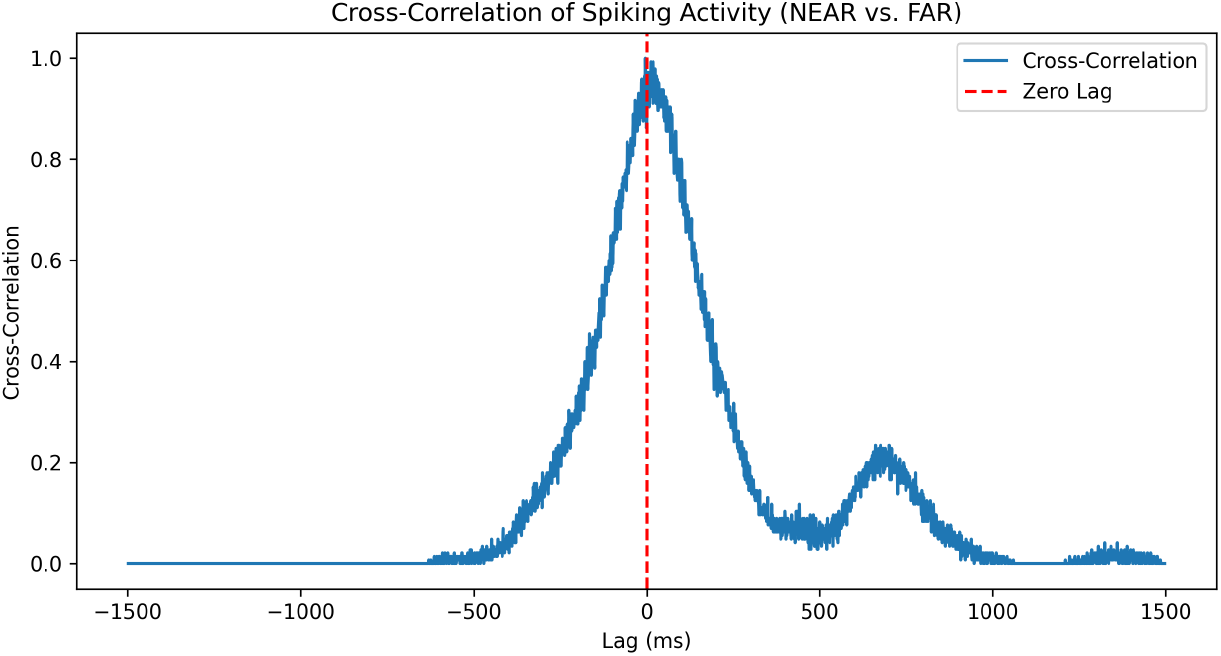
Cross-correlation of spiking activity between NEAR and FAR conditions. The peak at zero lag indicates strong neural synchrony in the NEAR condition.

Conversely, the FAR condition showed reduced synchrony, as evidenced by a lower peak in the cross-correlation function. This indicates that neural activity in the frontal and occipital regions was less temporally coordinated when the flankers were farther away. These results suggest that the spatial proximity of distractors influences the degree of coordination between different brain regions involved in task performance, with closer distractors requiring more synchronized neural processing.

## 4 Discussion

The results of this study provide strong evidence that the spatial proximity of distractor stimuli significantly modulates neural processing during visual tasks. The increased low-frequency power (delta and theta) and elevated firing rates observed in the NEAR condition suggest that when flankers are positioned closer to the target stimulus, the brain experiences heightened cognitive load and neural engagement. This finding is consistent with prior research on the flanker effect, which has demonstrated that closer distractors increase interference, necessitating greater attentional control and cognitive resources.

A central finding of this study is the difference in spiking activity between the NEAR and FAR conditions. As illustrated in Figure 10, the NEAR condition exhibits a higher frequency of spiking events, with dense clusters of neural firings occurring over time. This increased activity suggests that closer flankers demand greater neural engagement, likely due to heightened inhibitory control mechanisms required to suppress interference. In contrast, the FAR condition shows more dispersed and less frequent spiking, indicating that the brain processes the target stimulus with greater efficiency when distractors are positioned further away.

**Fig. 10.**
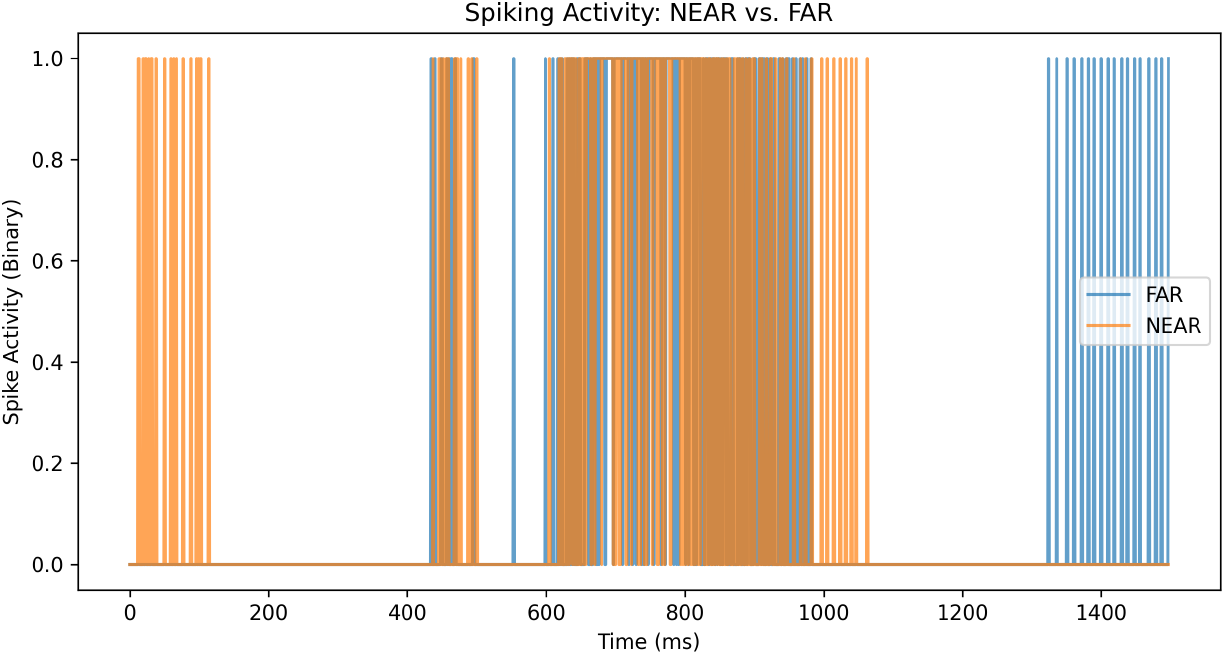
Spiking activity for NEAR and FAR conditions. The x-axis represents time in milliseconds, while the y-axis represents binary spike activity. The NEAR condition (orange) exhibits more frequent and clustered spiking activity compared to the FAR condition (blue), indicating higher neural engagement when distractors are closer to the target stimulus.

Further supporting this finding, the distribution of spike times presented in Figure 11 highlights the variability in neural responses under both conditions. The violin plot demonstrates that the NEAR condition exhibits a broader and more irregular distribution of spike times, suggesting increased neural variability and sustained activation. This result aligns with the interpretation that closer distractors generate greater cognitive load, leading to heightened but inconsistent neural firing patterns. Conversely, the FAR condition presents a more concentrated distribution of spike times, implying more stable and efficient neural processing with reduced interference.

**Fig. 11.**
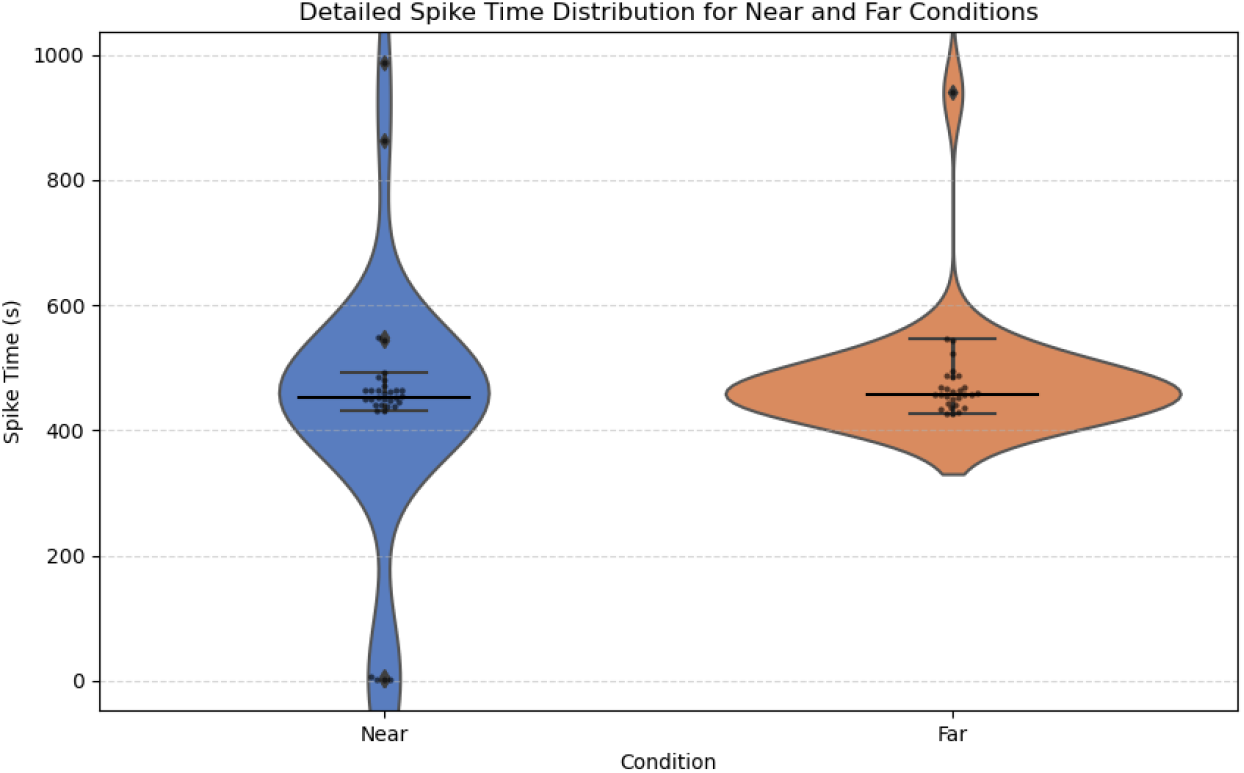
Violin plot showing the distribution of spike times for NEAR and FAR conditions. The NEAR condition (blue) exhibits a broader distribution with higher variability, indicating irregular neural firing patterns. In contrast, the FAR condition (orange) shows a more concentrated distribution, suggesting more stable and less frequent spike activity.

A quantitative measure of the relationship between the NEAR and FAR conditions is provided by the correlation analysis of spiking activity. The computed correlation coefficient of *r* = 0.4283 indicates a moderate positive relationship between the two conditions, suggesting that while neural firing patterns in NEAR and FAR conditions share some similarities, substantial differences persist. This moderate correlation reinforces the interpretation that spatial proximity of flankers induces distinguishable neural processing patterns, with NEAR conditions exhibiting more pronounced variability and intensity of neural activity.

These findings provide further support for the hypothesis that the NEAR condition imposes greater cognitive demands. The elevated firing rates observed in this condition likely reflect the increased burden on neural circuits associated with attentional control and inhibitory processes. This finding aligns with theoretical models suggesting that the spatial proximity of distractors affects neural engagement, with closer distractors requiring greater effort to filter out irrelevant stimuli.

## 4 Conclusion

In this study, we conducted a comprehensive multimodal investigation of neural responses to visual stimuli in near and far flanker conditions. By analyzing EEG event data across multiple subjects, we examined key neural activity markers, including spectrograms, cross-correlation, wavelet transformations, and time-frequency decompositions across alpha, beta, theta, gamma, and delta bands. Additionally, estimated neuron firing rates, spike activity distributions, and comparative statistical visualizations were utilized to discern meaningful differences between the two conditions.

Our findings indicate that neural responses exhibit condition-dependent variations, with distinguishable firing rate patterns and spectral signatures between the near and far flanker stimuli. These results support the hypothesis that the spatial configuration of visual stimuli influences neuronal processing dynamics. The statistical analysis of estimated firing rates further reinforces the observed differences, providing evidence of condition-specific neural engagement.

This research contributes to the broader understanding of neural mechanisms underlying visual perception and cognitive processing in competitive attentional environments. Future work will focus on refining computational models to enhance the interpretability of neural activity, integrating advanced machine learning techniques, and expanding the dataset to explore inter-subject variability more extensively.

## References

[1] Association, B.N., et al.: Neuroscience: Science of the brain-an introduction for young students. Liverpool: British Neuroscience Association (2003)

[2] Herrero, J.L.: Neurophysiology and neuropharmacology of visual attention. PhD thesis, Newcastle University (2011)

[3] Léveillé, J., Versace, M., Grossberg, S.: Spiking dynamics during perceptual grouping in the laminar circuits of visual cortex. Technical report, Boston University Center for Adaptive Systems and Department of Cognitive … (2009)

[4] Léveillé, J., Versace, M., Grossberg, S.: Running as fast as it can: How spiking dynamics form object groupings in the laminar circuits of visual cortex. Journal of Computational Neuroscience 28(2), 323–346 (2010)

[5] Nigbur, R., Ivanova, G., Stürmer, B.: Theta power as a marker for cognitive interference. Clinical Neurophysiology 122(11), 2185–2194 (2011)

[6] Szczepanski, S.M., Crone, N.E., Kuperman, R.A., Auguste, K.I., Parvizi, J., Knight, R.T.: Dynamic changes in phase-amplitude coupling facilitate spatial attention control in fronto-parietal cortex. PLoS biology 12(8), 1001936 (2014)

[7] Terhune-Cotter, B., Holcomb, P.J., Midgley, K.J., Ortega, S.E., Akers, E.M., Emmorey, K.: “Flankers-FAR”. 10.18112/openneuro.ds005868.v1.0.1

[8] Terhune-Cotter, B., Holcomb, P.J., Midgley, K.J., Ortega, S.E., Akers, E.M., Emmorey, K.: “Flankers-NEAR”. 10.18112/openneuro.ds005866.v1.0.1

[9] Anderson, B., Sheinberg, D.L.: Effects of temporal context and temporal 13 expectancy on neural activity in inferior temporal cortex. Neuropsychologia 46(4), 947–957 (2008)

[10] Berlemont, K., Nadal, J.-P.: Perceptual decision-making: Biases in post-error reaction times explained by attractor network dynamics. Journal of Neuroscience 39(5), 833–853 (2019)

[11] Cohen, B.P., Chow, C.C., Vattikuti, S.: Dynamical modeling of multi-scale variability in neuronal competition. Communications Biology 2(1), 319 (2019)

[12] Mallot, H.A.: Computational Neuroscience vol. 486. Springer, ??? (2013)

[13] Thut, G., Miniussi, C., Gross, J.: The functional importance of rhythmic activity in the brain. Current Biology 22(16), 658–663 (2012)

[14] Yang, X., Fiebelkorn, I.C., Jensen, O., Knight, R.T., Kastner, S.: Differential neural mechanisms underlie cortical gating of visual spatial attention mediated by alpha-band oscillations. Proceedings of the National Academy of Sciences 121(45), 2313304121 (2024)

[15] Yang, X.: Attention in space and time: From behavior to neural mechanisms. PhD thesis, Princeton University (2023)

[16] Marquardt, L.A.: Eeg analysis in adults with attention-deficit-hyperactivity-disorder. resting state and behavioural data analysis. Master’s thesis, The University of Bergen (2015)

